# Mitochondrial structure despite nuclear panmixia: sex-specific dispersal dictates population structure in sperm whales

**DOI:** 10.1101/2025.04.29.651281

**Authors:** Reid S. Brennan, Lynsey A. Wilcox Talbot, Anthony Martinez, Lance P. Garrison, Dan Engelhaupt, Nicole L. Vollmer, Patricia E. Rosel

## Abstract

Marine mammals have high potential for dispersal, yet behavioral or environmental constraints can limit gene flow. This is true for the endangered sperm whale, *Physeter macrocephalus*, which has a global distribution and long-distance migrations. While previous studies revealed mitochondrial population structure with weak nuclear structure globally, genomic approaches examining this pattern have been limited. Understanding connectivity is critical for the management of this species due to population declines relative to pre-whaling, recent impacts from the *Deepwater Horizon* oil spill, and ongoing threats from anthropogenic sound. We investigated the connectivity between two regions, the U.S. Gulf of America (Mexico) and western North Atlantic Ocean, using reduced representation genomics and mitochondrial control region sequencing of 73 sperm whales. We found that relatedness decreased with geographic distance, likely due to the presence of familial structure. Nuclear markers showed no population structure (*F*_ST_ = 0.001-0.008), while mitochondrial structure was high (*F*_ST_ = 0.36-0.65), consistent with male-biased dispersal and female philopatry. Across all samples, genetic diversity (nuclear: 0.0014; mitochondrial: 0.0017) and effective population size (N_e_=460) were low. Given this low diversity and evidence for the partitioning of genetic variation, we recommend managers treat these two regions as distinct in order to preserve existing variation and promote resilience of this species. These results show that despite the increased power of a genomic approach, it is essential to consider the biology of the species at hand and leverage both mitochondrial and nuclear markers to understand the genetic structure of threatened species.

## Introduction

Marine environments are paradoxical: despite few obvious physical barriers to gene flow and dispersal (Palumbi, 1994; Grosberg and Cunningham, 2001), many species show population structure and genetic differentiation (Kelly and Palumbi, 2010; Sanford and Kelly, 2011). While population structure may be expected for organisms with low dispersal potential and restricted or patchy habitat use (Bohonak, 1999), this pattern is seen even for species with extremely high dispersal potential (Ansmann *et al*., 2012; Mamoozadeh *et al*., 2020). Understanding these patterns of population structure is critical, particularly for species of conservation or economic concern, where knowledge of population connectivity and structure directly informs management decisions, such as the strategic design of marine reserves (Palumbi, 2003; Cros *et al*., 2017; Bernatchez *et al*., 2017).

For species with a passive pelagic life stage, realized dispersal is highly dependent on local currents that can either restrict or extend dispersal range (Largier, 2003; Byers and Pringle, 2006; Morgan *et al*., 2009; Álvarez-Noriega *et al*., 2020). However, successful gene flow requires not only physical movement, but also survival in the new habitat; evidence suggests that post-settlement mortality may be higher in long-distance dispersers (Burgess *et al*., 2012). In contrast, species with the ability to swim are affected by different processes such as philopatry, where individuals return to reproduce in their natal regions, which can greatly limit gene flow (Hendry *et al*., 2004). In extreme cases such as sea turtles, philopatry and natal homing can drive population structure despite their long distance migration and dispersal (Komoroske *et al*., 2017). In contrast, a combination of local adaptation and genetic drift has resulted in the emergence of sympatric ecotypes of killer whales (*Orcinus orca*) that are differentiated by prey type (Moura *et al*., 2014). Thus, despite the potential for high gene flow, genetic differentiation is common in oceans and driven by multiple mechanisms.

Understanding population connectivity and structure is particularly crucial for broadly distributed marine mammals with high dispersal potential, as this knowledge directly informs effective management strategies. The sperm whale, *Physeter macrocephalus*, exemplifies this need as their global distribution and dominant role as predators in the deep sea make them critical to ecosystem functioning (Whitehead, 2018). For example, sperm whales influence nutrient cycles by transporting deep-sea nutrients to the photic zone via defecation, which stimulates primary production and is an important component of the biological carbon pump and carbon export to deep waters (Lavery *et al*., 2010). Therefore, conservation of this species is critical to maintain ecosystem functioning of, and carbon sequestration in, our deep oceans.

Sperm whales exhibit social structure where females are philopatric and maintain long-term familial social groups while males are highly dispersive and relatively solitary after maturity, except during breeding (Whitehead, 1993; Richard *et al*., 1996). This sex-biased dispersal pattern is reflected in their genetic structure: mitochondrial markers reveal clear population differentiation across ocean basins (Alexander *et al*., 2016), while nuclear markers often find weaker or no evidence of structure (Engelhaupt *et al*., 2009; Mesnick *et al*., 2011a). Whaling has impacted populations, with historical worldwide population sizes estimated to be around 2 million, but 2022 population sizes of about 845,000; there has been an international moratorium on commercial whaling since 1986 (Whitehead and Shin, 2022). As such, the species is currently listed as endangered under the U.S. Endangered Species Act and vulnerable on the International Union for Conservation of Nature (IUCN) (2024) Red List. While mitochondrial genetic diversity is low globally (Alexander *et al*., 2013), this pattern appears to predate whaling impacts and was likely driven by a species-wide bottleneck approximately 125 thousand years ago, followed by a population expansion over the last 20-40 thousand years (Morin *et al*., 2018).

While there has been a global focus on understanding sperm whale population structure, the connectivity between individuals from the U.S. Gulf of America (formerly the U.S. Gulf of Mexico; hereafter “Gulf”) and Atlantic Ocean remains poorly understood. This is critical as the species is susceptible to habitat threats including anthropogenic sound and large-scale environmental disasters. For example anthropogenic sound stems from seismic operations related to oil and gas activity as well as high ship traffic (Estabrook *et al*., 2016) and it is estimated that 16.1% of the sperm whales in the Gulf were exposed to oil from the *Deepwater Horizon* spill (Solsona-Berga *et al*., 2024); it is likely that both of these factors have led to local population declines. Namely, the *Deepwater Horizon* oil spill was the primary driver of increased strandings of marine mammals during an unusual mortality event following the spill (Deepwater Horizon Natural Resource Damage Assessment Trustees, 2016). Understanding population connectivity between the Gulf and other regions is essential for evaluating whether they should be managed as distinct units and for assessing their resilience and response to region-specific threats. Previous genetic analyses using nuclear microsatellites and low resolution, single nucleotide polymorphism (SNP) based analyses have shown, in some cases, limited power to detect population differentiation in this species (Engelhaupt *et al*., 2009; Mesnick *et al*., 2011b; Alexander *et al*., 2016), and genome-wide approaches have only been applied to a few regions (i.e., Violi et al., 2023). Here, we fill this gap by using a reduced representation approach to characterize the genome-wide genetic variation of 73 sperm whale samples collected from 1995 to 2014 across the northern Gulf and western North Atlantic. By combining these genomic data with additional data from the mitochondrial control region, we reveal the genetic relationships between geographic regions, relatedness between individuals, and the standing levels of genetic variation; factors that are essential for long-term resilience to changing global conditions. These results provide crucial information for developing conservation strategies for this vulnerable species across its U.S. range.

## Methods

Skin samples were collected from sperm whales in the northern Gulf and western North Atlantic from 1995 to 2014 and consisted of a mix of stranded individuals (n=11) and remote biopsies (Sinclair *et al*., 2015) from free swimming animals (n=62), resulting in 73 total samples (Figure 1, Supplemental Metadata). Note that all 11 stranded individuals were from the Atlantic coast. DNA was extracted from skin using a Qiagen DNeasy Blood and Tissue kit following the manufacturer’s protocol and four samples with brown-colored elutions were further purified with MO BIO’s PowerClean DNA Clean-Up Kit (Qiagen) to remove potential PCR inhibitors. DNA quality was determined by gel electrophoresis and quantity was read on a fluorometer (GE Healthcare Hoefer DyNA Quant 200 Fluorometer). A subset of the biopsy samples (n=40) were previously sexed (Engelhaupt *et al*., 2009) and the remaining were genetically determined by PCR amplification of ZFX and SRY gene fragments as described by Rosel (2003), except 25 ng of DNA and 0.75 U of *Taq* DNA Polymerase (Invitrogen) were used in each reaction. PCR fragments to determine sex were visualized by gel electrophoresis on a 2.5% agarose gel in 1x SB (sodium borate) buffer. The sex of stranded individuals was determined in the field through visual inspection. Samples were separated into four putative geographic regions based on their sampling location and hypothesized population structure and separation: Atlantic, Dry Tortugas, Northern Gulf (N. Gulf), and Western Gulf (W. Gulf) (Fig. 1).

**Figure 1:**
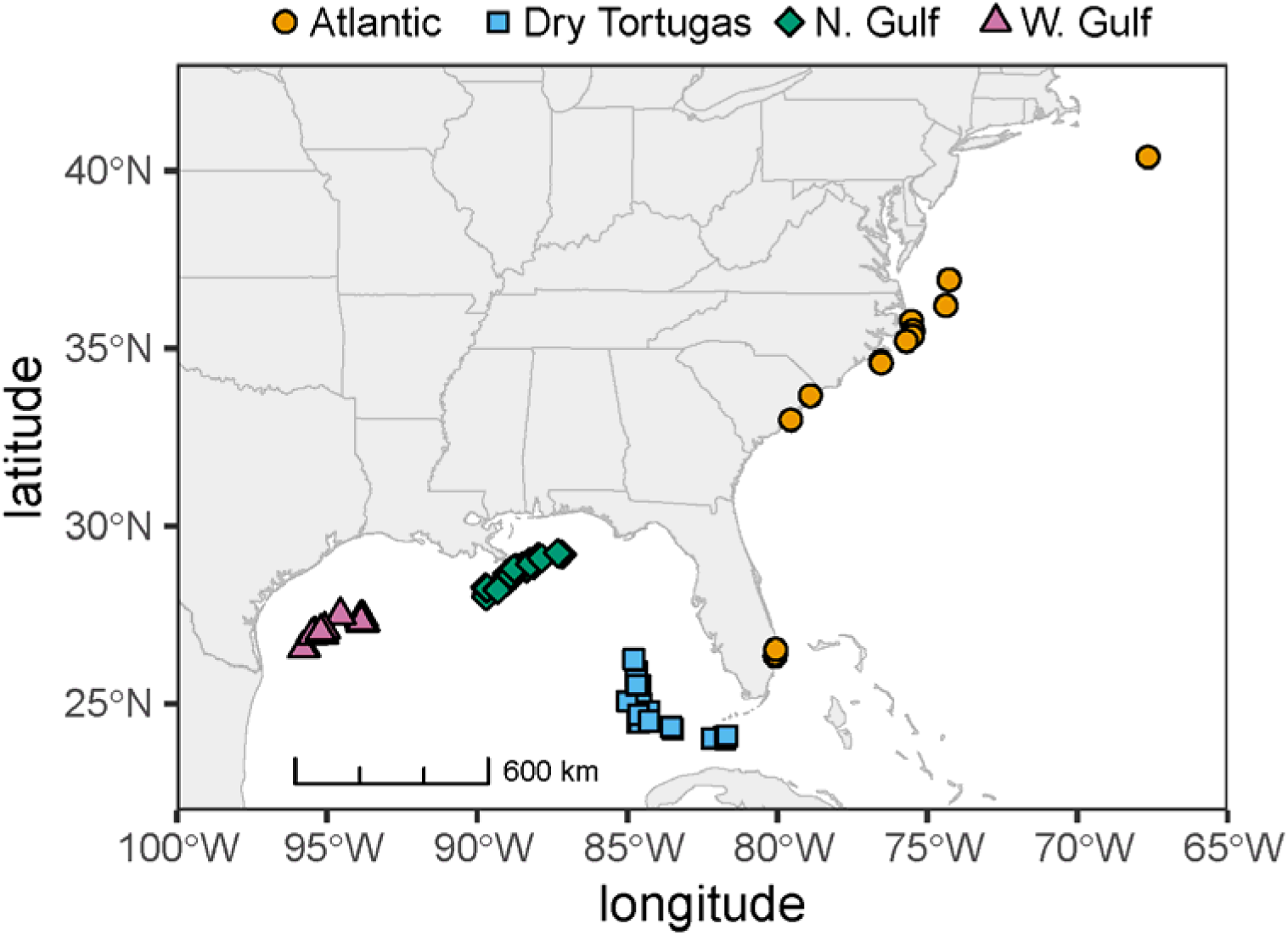
Collection sites for all samples. Each point is an individual sample and points are colored by geographic region corresponding to the *a priori* hypothesized population structure.

Genomic data were collected using a Nextera-tagmented reductively-amplified DNA (nextRAD) genotyping-by-sequencing approach from SNPsaurus (LLC). This method is a reduced representation approach that uses selective PCR primers to repeatedly target the same genomic regions across samples. Genomic DNA was randomly fragmented with the Nextera reagent (Illumina) and then amplified with a custom primer targeting the 10 base pair (bp) sequence GTGTAGAGCC. Resulting libraries were sequenced on one lane of an Illumina HiSeq 4000 with 150 bp single end reads. See Russello et al., for full details of nextRAD (2015).

Raw reads were quality checked with fastQC v0.11.9, trimmed for quality and adapter contamination using *fastp* v0.23.4 (Chen *et al*., 2018), and aligned to the *Physeter macrocephalus* reference genome (Fan *et al*., 2019) using bwa-mem2 v2.2.1 (Vasimuddin *et al*., 2019). Variants were called with freebayes v1.3.6 (Garrison and Marth, 2012) and stringently filtered using VCFtools v0.1.16 (Danecek *et al*., 2011) to keep only sites that were: bi-allelic, minor allele frequency greater than 0.05, less than 30% missing data, and a minimum mean depth of 10 and maximum of 53. Finally, potential paralogs were removed using HDplot (McKinney *et al*., 2017). This resulted in 4,441 single nucleotide polymorphisms (SNPs). See accompanying scripts on Zenodo for full filtering details.

Related individuals were identified using ngsRelateV2 (Hanghøj *et al*., 2019) using both the KING-robust kinship estimator (Waples *et al*., 2019) and the coefficient of coancestry Θ (Jacquard, 1974). Individuals were considered related if they were at least 3rd-degree relatives, with a kinship coefficient for KING greater than 0.0442 (Manichaikul *et al*., 2010) and Θ greater than 0.0625 (Jacquard, 1972). We also took a more lenient approach, considering individuals related if they had either a kinship coefficient greater than 0.05 or Θ greater than 0.05.

Individuals with Θ of ∼0.25 were considered full-siblings and 0.125 half-siblings. We tested for a relationship between relatedness (Θ) and ocean distance between samples using a zero-inflated beta regression using the GAMLSS framework (Rigby and Stasinopoulos, 2005) in R v4.4.1 (R Core Team, 2024). This approach was chosen as Θ is bounded between zero and one and contains zeros. Pairwise ocean distances between all individuals were calculated by finding the least-cost path using the marmap package (Pante and Simon-Bouhet, 2013), limiting minimum depth to 0.3 meters to avoid crossing land; paths were visually checked to ensure land was not crossed. While some individuals were strandings, these location estimations represent an approximate location of the individuals prior to stranding, though the distance any individual drifted prior to stranding cannot be known.

To understand the potential genetic structure among geographic regions, we first conducted a principal component analysis (PCA). For all analyses of population structure, individuals were removed such that no related individuals remained in the dataset, leaving 61 unrelated individuals. Using PLINK v1.90, variants were thinned for linkage disequilibrium (LD) with the option –indep 50 5 2 and a PCA was run on the resulting 3,977 SNPs. Ancestries of each individual were estimated using Admixture v1.3.0 (Alexander *et al*., 2009), testing one to eight populations, and the most likely number of populations was determined based on the lowest cross validation error. We repeated these same analyses with only the 42 female individuals. Pairwise *F*_ST_ among geographic regions was calculated using Weir and Cockerham’s method (Weir and Cockerham, 1984) implemented in snpR v1.2.11 with 500 bootstrap permutation replicates to calculate significance (Hemstrom and Jones, 2023).

Population structure was also tested for using discriminant analysis of principal components (DAPC). One of the most important parameters in running DAPC is the number of principal component (PC) axes to include as the predictor. Thia (2023) recently showed that the number of PCs should be equal to *k*-1 where *k* is the number of effective populations in the dataset (Thia, 2023). We applied this principle as well as the cross-validation and *a*-score approach described by the authors of the adegenet package (Jombart, 2008). The number of populations was estimated using K-means clustering (*find.clusters()*) where the most likely *k* was identified as the lowest Bayesian information criterion (BIC) score and a system with population structure should give an “elbow” where BIC values decrease until the most likely *k* and then plateau or increase. This pattern should be robust to changes in the number of PCs used when structure is present. Following this, we tested the hypotheses that the number of populations is equal to four, the number of geographic locations, or two, the split between Gulf and Atlantic individuals. We assessed the fit of each of these models using 100 replicates of training-testing partitioning as recommended by Thia (2023). For each replicate, we randomly assigned 70% of the samples to the training partition and built a DAPC model. We then predicted the population source of the other 30% of individuals (testing partition) given this model. Assignment accuracy was calculated as the proportion of testing individuals assigned back to their population of origin. With a two population model, random assignment from an uninformative model would result in 50% accuracy while a four population model would result in 25% accuracy.

Across all calculations of genetic diversity and inbreeding, potential siblings were retained in the dataset to avoid inducing bias (Waples and Anderson, 2017). All values were calculated by the putative geographic population as well across all individuals. Pairwise genetic diversity (π) was estimated in 50 kb windows across assembled chromosomes (and not unassembled scaffolds) using *pixy* v1.2.10 (Korunes and Samuk, 2021). Both variant and invariant sites were called with freebayes v1.3.6 and filtered by depth and missingness in the same manner as described above. Including invariant sites is critical to avoid bias in estimates of π. Confidence intervals were determined using 10,000 bootstrap replicates and results were robust when window sizes were decreased to 10 kb. *F*_IS_ and expected and observed heterozygosity were calculated using Stacks v2.65 (Rochette *et al*., 2019). Contemporary effective population size was estimated using *currentNe* v1.0 (Santiago *et al*., 2024), using only loci located on assembled chromosomes and specifying the average number of siblings an individual has in the sample (0.11), as well as with NeEstimator v2 (Do *et al*., 2014b) implemented via dartR v2.9.8 (Mijangos *et al*., 2022).

To quantify mitochondrial genetic diversity and population structure, we sequenced the mitochondrial DNA (mtDNA) control region for all 73 samples. The control region was PCR amplified using primers L15928 (Vollmer *et al*., 2011) and a modified version of H00651 (5′- CAGAAGGCTAGGACCAAACCT-3′, modified from Kocher (1989)) and Phusion^TM^ High-Fidelity DNA Polymerase (Thermo Scientific). Each PCR was performed in a 25 µL reaction with 1 x Phusion^TM^ HF Buffer, 0.5 U Phusion^TM^ Polymerase, 0.5 µM of each primer, 200 µM dNTPs, and 25 ng of DNA. The thermal cycling profile consisted of an initial denaturation step of 98°C for 30 s followed by 30 cycles of 98°C for 10 s, 63°C for 20 s, and 72°C for 20 s with a final extension at 72°C for 5 min. Amplified products were purified via extraction from low melting point agarose gels followed by digestion using agarase (Sigma-Aldrich) and then sequenced in both directions using a BigDye Terminator v3.1 Cycle Sequencing Kit (Thermo Scientific) on an ABI 3130 Genetic Analyzer. The forward and reverse sequences were independently edited and then consensus sequences were assembled using Sequencher v5.3 (GeneCodes). Sequences were aligned using Se-Al 2.0a11 (Rambaut, 2002) and flanking regions were truncated to include only the control region sequence of 955 bp. Nucleotide diversity was calculated for all samples together and as separate populations using Pegas v1.3 (Paradis, 2010). Hierfstat v0.5.11 was used to calculate pairwise *F*_ST_ between populations with 1000 bootstrap replicates to determine the 95% confidence intervals (Goudet, 2005). Statistical parsimony haplotype networks (i.e., TCS for Templeton, Crandall and Sing (1992)) were calculated using PopART (Leigh *et al*., 2015). Finally, we ran an Analysis of Molecular Variance (AMOVA) using ade4 v1.7.22 (Dray and Dufour, 2007) implemented in poppr v2.9.6 (Kamvar *et al*., 2014).

## Results

### Relatedness

There were four pairs of first-degree relatives such as full siblings or parent offspring (seven females, one male), eight pairs of second-degree relatives (eight females, six males, two unknown), and four pairs of third-degree relatives (Fig. 2; seven females, one male); the number of third-degree relatives should be treated cautiously as these relationships can be difficult to identify (Manichaikul *et al*., 2010). Full-siblings were sampled within a geographic region, where 3 pairs were sampled within the W. Gulf and one within the N. Gulf (Fig. 2, S1). Overall, closely related individuals were more likely to be sampled at closer geographic distances, with relatedness (Θ) decreasing by 0.036 per 1000 km (p = 0.003), likely reflecting the familial structure of sperm whales (see discussion).

**Figure 2:**
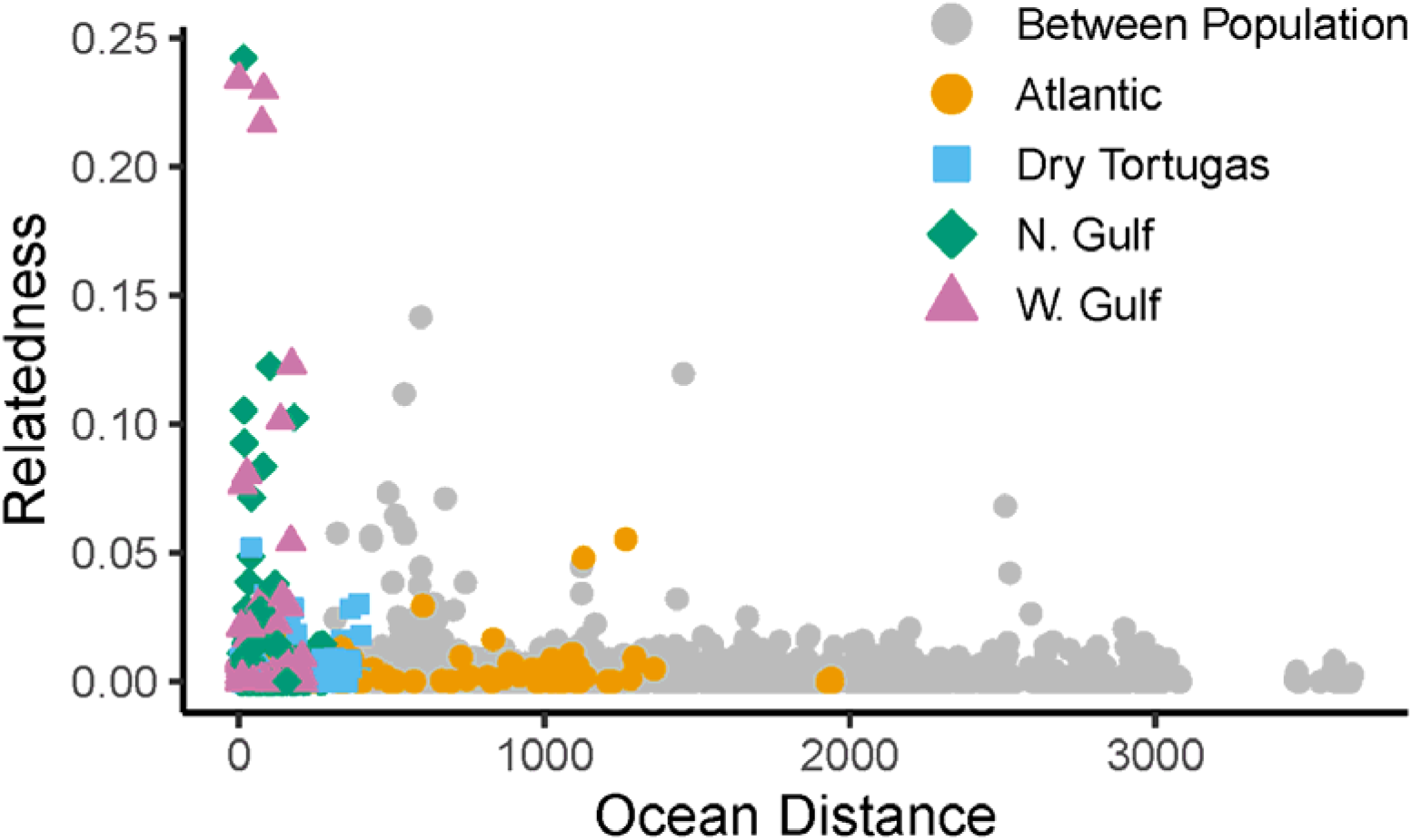
Pairwise relatedness between all individuals colored by population origin of each individual. Relatedness is Θ and distance is the shortest ocean distance in km, avoiding land between two individuals. There was a negative relationship between ocean distance and relatedness where individuals collected at close ocean distance were more likely to be closely related than those at farther distances (p = 0.003).

### Nuclear DNA structure and diversity

There was no evidence for population structure when looking at the nuclear nextRAD markers. Principal component analysis showed panmixia and no clustering by population (Fig. 3A, S2). Similarly, *F*_ST_ was nearly zero between all populations (*F*_ST_=0.001-0.008; Fig. 3B, above diagonal) and not significantly different from the null hypothesis of panmixia (Fig. 3B, below diagonal). The most likely number of populations was one, as indicated by the cross-validation error from Admixture (Fig. S3). Finally, DAPC revealed no power to discriminate among populations using either a two- or four-population model and regardless of the approach for the number of PCs included (Fig. 3C; S4). The training-testing partitioning was no better than random in assigning individuals to population, indicating a lack of discriminatory genetic differences among populations (Fig. 3C). This result was robust to the number of PCs included in the DAPC model (Fig. S4). Further, K-means clustering showed no “elbow” indicative of population structure (Fig. S5) and when adding PCs to the DAPC model the slope of the curve switched from negative (low PCs) to positive (high PCs), mirroring simulation results from Thia (2023) showing the same pattern in panmictic populations; systems with population structure are generally robust to changes in the number of PCs included in the model. Note that the lack of population structure is consistent when analyzing the unrelated females without any males (Fig. S6, S7).

**Figure 3:**
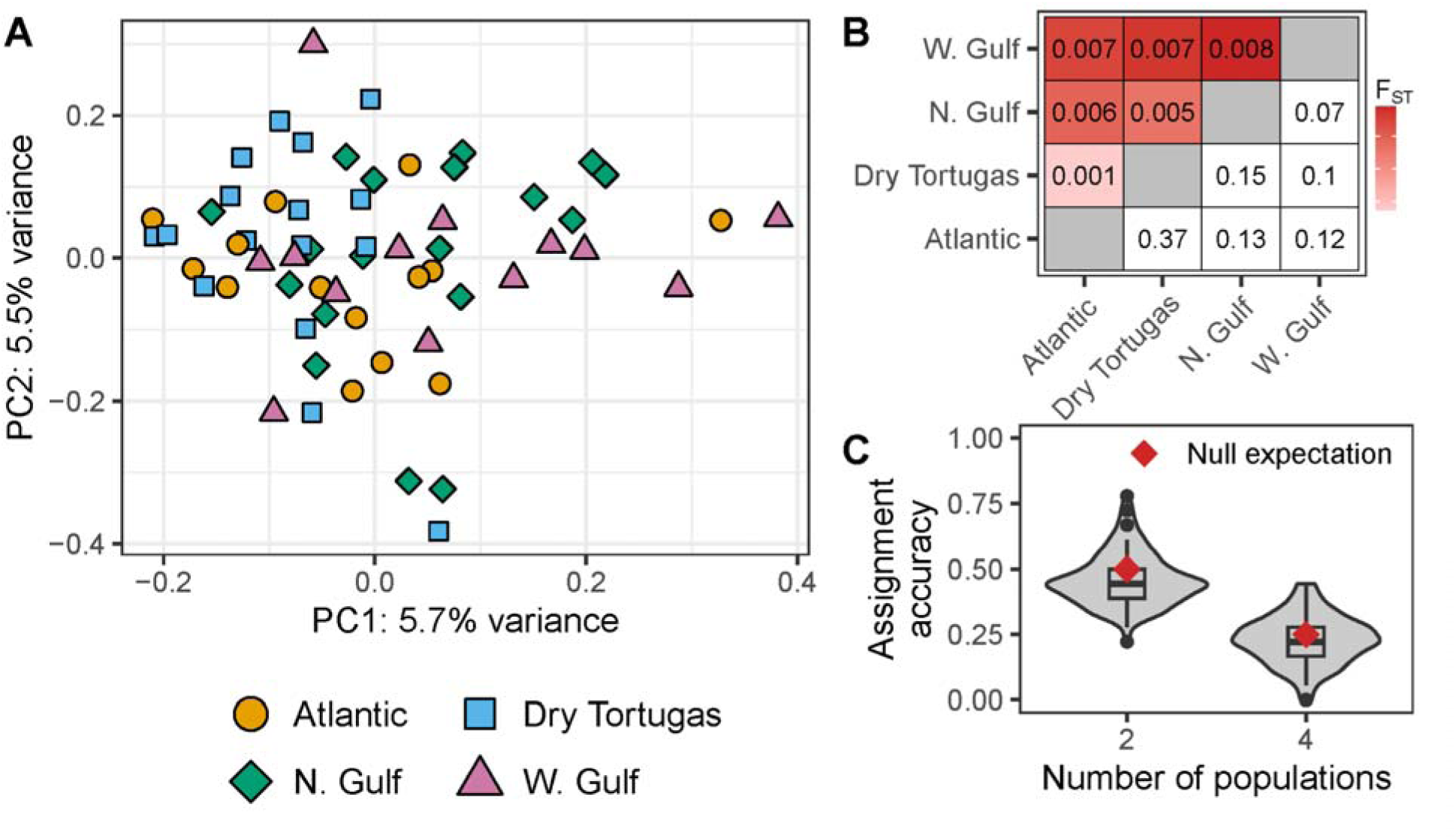
Lack of population structure with nuclear DNA indicates panmixia across populations. A. Principal components analysis of 3,977 LD thinned SNPs showing lack of clustering by geographic location. Geographic locations are represented by shape and color. B. Pairwise *F*_ST_ estimates between locations. Above diagonal and colors indicate *F*_ST_ values while below the diagonal represents p-values from 500 bootstrap permutations testing panmixia. C. Assignment accuracy from DAPC following 100 replicates of training-testing partitioning assuming both two (Atlantic: Atlantic + Dry Tortugas; Gulf: N. Gulf + W. Gulf) and four populations. If population assignments were completely random, we expect 50% accuracy for the two-population model and 25% accuracy for the four-population model. The boxplot and violin plot show the distribution of mean accuracy for the 100 replicate runs and the red diamond is the null expectation given random assignments.

For the nuclear markers, across all individuals, pairwise genetic diversity (Tajima’s π) was 0.00136 (95% confidence interval: 0.00133, 0.00140; Fig. 4A). As there was no evidence of population structure, all individuals should probably be treated as a single population for these calculations. For completeness we also provide location-level estimates: Atlantic: 0.00137 [0.00133, 0.00141]; Dry Tortugas: 0.00133 [0.00130, 0.00137]; N. Gulf: 0.00140 [0.00136, 0.00143]; W, Gulf: 0.00132 [0.00128, 0.00136]. Only N. Gulf and W. Gulf do not overlap in their confidence intervals. Observed heterozygosity was slightly lower than expected heterozygosity across all individuals (H_obs_: 0.263 [0.261, 0.265]; H_exp_: 0.283 [0.280, 0.284]; Fig. 4B,C).

**Figure 4:**
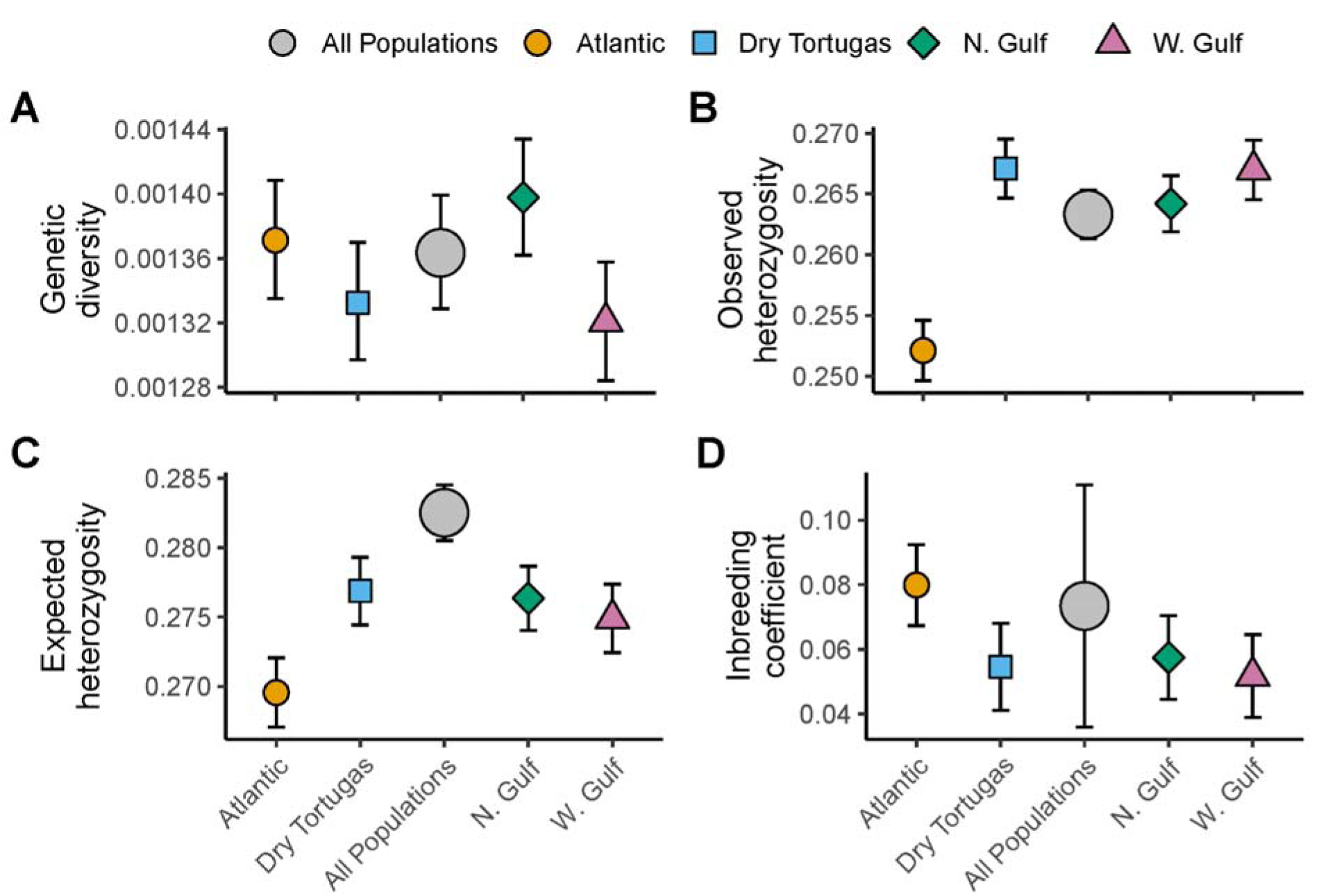
Measures genetic diversity for nuclear markers: pairwise genetic diversity (A), observed (B) and expected (C) heterozygosity, and inbreeding coefficients (D; *F*_IS_). Values were estimated considering all individuals as a single population (“All populations”), as suggested by population structure results, as well as by geographic location.

However, estimates were largely consistent across location, with the exception of Atlantic, which was lower than the others(H_obs_: 0.252 [0.250, 0.255]; H_exp_: 0.270 [0.267,0.272]). The inbreeding coefficient (*F*_IS_) was 0.073 and did not overlap with zero [0.036, 0.111]. Again, there were few differences across locations except that Atlantic (0.0799 [0.067, 0.092]) was higher than W. Gulf (0.052 [0.039, 0.065]). Contemporary effective population size estimated across all geographic locations using CurrentNe was 459.54 (90% CI: 324.57, 650.64) without specifying the number of siblings in the sample and 708.52 (90% CI: 469.47, 1069.29) when specifying the average number of siblings in the sample. NeEstimator provided similar results (516.6) but with infinite confidence intervals, likely due to the sampling error correction (Do *et al*., 2014a).

### Mitochondrial DNA structure and diversity

Similar to the nuclear markers, there was low overall mitochondrial genetic diversity (0.0017 [0.0014, 0.0019]) and no significant difference in levels of diversity among locations(Fig. 5A). However, in contrast to the nuclear DNA there was significant evidence for population structure. The haplotype network (Fig. 5B) showed segregation of haplotypes between locations and the AMOVA indicated significant population structure with 41.38% of the genetic variation occurring between locations and 58.62% occurring within locations (p = 0.01). *F*_ST_ between Atlantic and Dry Tortugas as well as between N. Gulf and W. Gulf were low and not significantly different from the null hypothesis of panmixia (Fig. 5C; Atlantic vs. Dry Tortugas: *F*_ST_ = 0.01, p=0.29; N. Gulf vs. W. Gulf: *F*_ST_ = 0.00, p = 1.00). In contrast, *F*_ST_ was high between Atlantic or Dry Tortugas versus N. Gulf or W. Gulf (p = 0.001; Fig. 5C). Together, these results indicate significant population structure for mtDNA between the Gulf versus Dry Tortugas and Atlantic.

**Figure 5:**
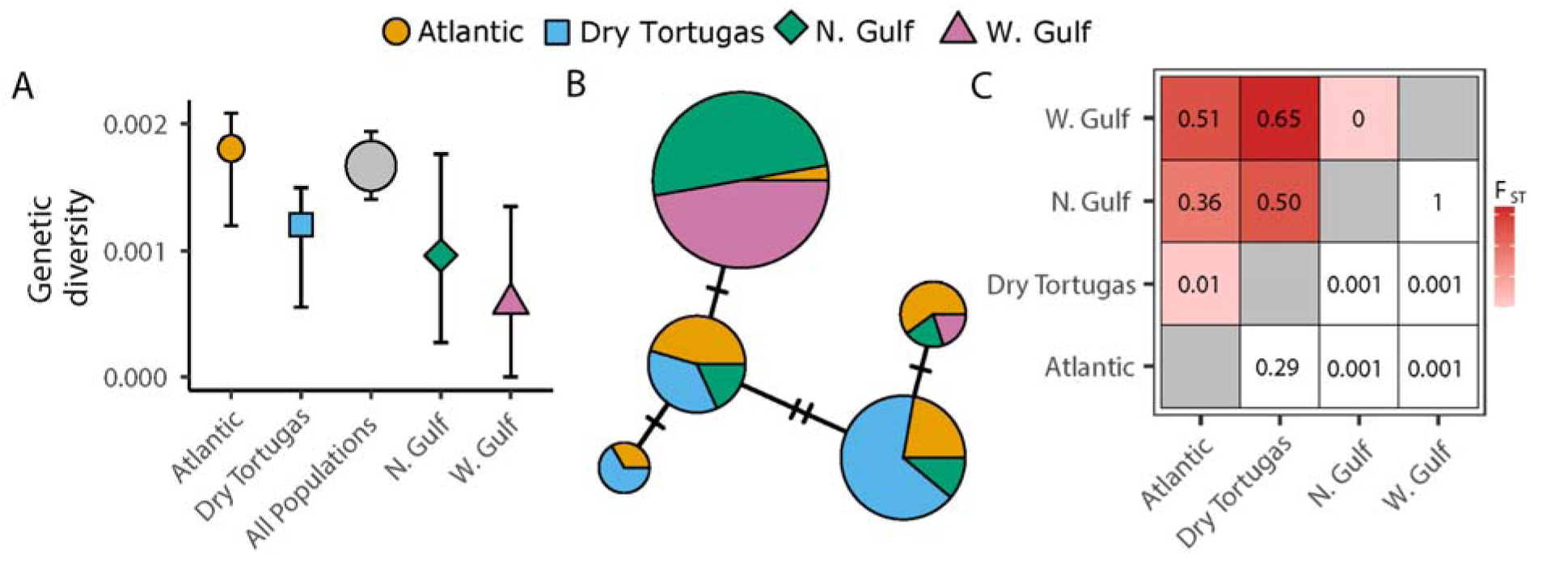
mtDNA population structure and diversity. A. Estimates of nucleotide diversity with 95% confidence intervals. B. Haplotype network calculated via the TCS method where the size of the circle indicates the number of individuals possessing each haplotype and color indicates the population origin. Hash marks denote the number of mutations between haplotypes. C. Measures of pairwise *F*_ST_ between populations where the above diagonal red triangle indicates *F*_ST_ values and the lower diagonal includes p-values from 1000 bootstrap permutations testing the null hypothesis of panmixia.

## Discussion

Here, we demonstrate that the highly dispersive sperm whale, *Physeter macrocephalus*, shows significant population structure in its mitochondrial DNA despite panmixia in its nuclear genome across the Northern Gulf and western North Atlantic Ocean. This result is contrary to the majority of population genetic studies where leveraging more variable markers, such as thousands of markers across the nuclear genome or microsatellites, is typically more powerful in revealing population structure than methods focused on single markers such as mitochondrial DNA (Gallego-García *et al*., 2021). This result is likely driven by the unique life history of sperm whales; there is strong family structure where females typically associate with their familial group for their entire lives and dispersal is predominantly male mediated (Best, 1979). Thus there is gene flow across the nuclear but not the mitochondrial genome. These results stress the importance of considering the life history of species and marker choice when assessing population structure, particularly for species such as sperm whales where management and conservation decisions may be based on the results. Further, our findings indicate that sperm whales in the Gulf are distinct from those in the Atlantic and should be considered separate evolutionary units for management purposes.

### Discrepancy between mitochondrial and nuclear population structure

In this study we adopted a reduced representation genomic approach to determine whether there was genetic differentiation between sperm whales from the Gulf and western North Atlantic. We hypothesized that the general lack of population structure for the nuclear genome in previous studies across various regions was due to a lack of power of microsatellite markers to detect fine-scale structure (Engelhaupt *et al*., 2009; Mesnick *et al*., 2011b; Alexander *et al*., 2016). Microsatellite loci are typically well suited to detecting neutral population structure due to their high mutation rate and allelic diversity and relatively low genotyping costs (Guichoux *et al*., 2011). However, microsatellites are usually limited to 10s of loci (Hodel *et al*., 2016) and under most scenarios, especially weak population differentiation, increasing the number of markers using genomic approaches can increase the power to detect this structure (Jeffries *et al*., 2016; Gallego-García *et al*., 2021; Poelstra *et al*., 2022). Similarly, increasing the number of loci can improve the ability to identify related individuals (Lemopoulos *et al*., 2019) and allows for accurate recovery of population structure and genetic diversity using fewer individual samples per population (Nazareno *et al*., 2017; Gallego-García *et al*., 2021). This final point is critical for sperm whales and other marine mammals where obtaining high sample numbers is particularly challenging and expensive.

Despite genotyping 4,441 nuclear loci across 73 samples, we found no evidence of population structure in the nuclear genome in this species between the Gulf and western North Atlantic, but evidence for mtDNA structure. The presence of mtDNA but not nuclear population structure is likely driven by the distinct sex-specific life history and dispersal of sperm whales; male-mediated dispersal and connectivity generate nuclear DNA panmixia but female philopatry leads to mtDNA structure. Sperm whales have multi-generational social structure and associations that consist of adult females and juveniles of both sexes. While males are found within these groups as juveniles, after reaching adolescence they disperse and become increasingly solitary with large ranges that extend to higher latitudes (Whitehead, 2018); males have been tracked moving between the Gulf and the Atlantic (Jochens *et al*.). The exception to this is during mating where males temporarily associate with females. In contrast, satellite tracking of sperm whales in the Gulf over weeks to almost two years has shown that females display strong philopatry and do not leave the region (OrtegaCOrtiz *et al*., 2012). Because mtDNA is inherited only matrilineally, the sex-specific dispersal of sperm whales can generate genetic differentiation at mitochondrial but not nuclear DNA, to which both males and females contribute. Previous work has found mtDNA structure between Gulf and Atlantic populations (Engelhaupt *et al*., 2009; Lizewski *et al*., 2025) as well as globally (Alexander et al., 2016; Morin et al., 2018), but very limited differentiation in the nuclear genome using microsatellite markers (Engelhaupt *et al*., 2009; Alexander *et al*., 2016). Similar patterns of mitochondrial but weak or missing nuclear population structure have been found in other species with similar life history strategies such as humpback whales (Baker *et al*., 2013), killer whales (Pilot *et al*., 2010), beluga whales (O’Corry-Crowe *et al*., 2018), and Port Jackson sharks (Day *et al*., 2019)

There was significant evidence that related individuals were more likely found at close geographic distances (Fig. 2), consistent with the life history of sperm whales. We found a mix of males and females within the closely related individuals (22 females, 8 males) and these individuals were generally within one geographic region. In this study, we do not know if the individual samples were taken from the same family group or the age of individuals. It is possible that the highly related individuals found here were members of the same group sampled at different times, which is particularly likely for the first degree relatives. Previous work has found that 82.5% of individuals within a social unit have a first degree relative in that unit, but has not found first degree relatives between social groups (Konrad *et al*., 2018). Our findings support this conclusion as the first degree relatives in this study were likely sampled from the same social groups with samples coming from the same day 0.5 km apart (Pmac097 and Pmac098; Θ = 0.23), six days and 16km apart (Pmac031 and Pmac125; Θ = 0.24), 3 days and 73 km apart (Pmac100 and Pmac102; Θ = 0.22), and 12 days and 89 km apart (Pmac093 and Pmac102; Θ = 0.23). The decay in relatedness with geographic distance likely reflects the lack of dispersal of females from their home range.

Nuclear genetic diversity (π) was 0.00136 (0.00133, 0.00140), which is on the lower end for mammals (Buffalo, 2021). Further, inbreeding was not high (Fig. 4D) and effective population size (*N_e_*) was ∼500 (459.54 [324.57, 650.64]). While this nearly satisfies the “50/500” rule where an *N_e_* of 50 is required to prevent inbreeding and 500 to maintain evolutionary potential and long-term viability (Franklin, 1980), more recent work shows that an *N_e_*of 1000 (Frankham *et al*., 2014) or even 5000 (Lande, 1995) is required for long-term viability of a population.

Therefore, *N_e_* is below what is considered resilient over the long term for this species. Given the substantial reductions in population numbers to about one third of pre-whaling sizes, it is key to continue to prioritize population recovery and protection to prevent further decreases in effective population size and long-term viability.

Finally, the mtDNA population structure and low genetic diversity presented here support the current delineation of the species into separate Northern Gulf and western North Atlantic management stocks (Hayes *et al*., 2021, 2024). While there is gene flow across the nuclear genome, the lack of mitochondrial gene flow indicates that sperm whales from the Gulf are evolutionarily distinct from the western North Atlantic, including the Dry Tortugas, and should be treated as separate populations. However, additional samples are needed to more accurately determine the geographic boundary between these populations; and to understand the wider connectivity among animals throughout the Gulf of America, Gulf of Mexico, and the Caribbean. Our estimate of mtDNA nucleotide diversity was 0.0017 [0.0014, 0.0019], which is slightly higher than the 0.001 estimate from Morin et al., (2018) based on 11 whole mitochondrial genomes.

This low mitochondrial diversity is likely driven by a bottleneck that occurred about 125 thousand years ago and an expansion over the past 20-40 thousand years (Morin *et al*., 2018). Given the low mitochondrial diversity in this species; preserving the current diversity across both the Gulf and Atlantic is likely key to maintaining the resilience of this species over the long-term.

## Acknowledgements

We thank Nicole Phillips for assistance with processing samples in the laboratory and Laura Dias, Jesse Wicker, and Wayne Hoggard for contributing to sample collection. Funding for sample collection was provided by the Bureau of Ocean Energy Management IA M09PG00014 and M11PG00041. This research was carried out, in part, under the auspices of the Cooperative Institute for Marine and Atmospheric Studies (CIMAS), a Cooperative Institute of the University of Miami and the National Oceanic and Atmospheric Administration, cooperative agreement #NA20OAR4320472. Biopsy sampling was conducted under NMFS MMPA permits 738, 779-1339, 779-1633, and 14450. The scientific results and conclusions, as well as any views or opinions expressed herein, are those of the author(s) and do not necessarily reflect those of NOAA or the Department of Commerce.

## Author Contributions

RSB conducted the analysis, made figures, and wrote the manuscript. LAWT conducted lab work and analyzed the mitochondrial data, while LAWT, NLV, and PER conceptualized the project and helped to interpret results. AM, LPG, and DE collected samples. All authors helped to revise the manuscript.

## Data Archiving

Raw genomic data can be found on NCBI under bioproject PRJNA1189584.

Code to run all analyses and metadata for samples, including NCBI accessions for mitochondrial data, can be found on Zenodo: https://doi.org/10.5281/zenodo.14715105

## References

IUCN. (2024). The IUCN Red List of Threatened Species. Version 2024–2. https://www.iucnredlist.org. Accessed on 2024-11-22.

Alexander DH, Novembre J, Lange K (2009). Fast model-based estimation of ancestry in unrelated individuals. Genome Res 19: 1655–1664.

Alexander A, Steel D, Hoekzema K, Mesnick SL, Engelhaupt D, Kerr I, et al. (2016). What influences the worldwide genetic structure of sperm whales (Physeter macrocephalus)? Mol Ecol 25: 2754–2772.

Alexander A, Steel D, Slikas B, Hoekzema K, Carraher C, Parks M, et al. (2013). Low Diversity in the Mitogenome of Sperm Whales Revealed by Next-Generation Sequencing. Genome Biol Evol 5: 113–129.

Álvarez-Noriega M, Burgess SC, Byers JE, Pringle JM, Wares JP, Marshall DJ (2020). Global biogeography of marine dispersal potential. Nat Ecol Evol 4: 1196–1203.

Ansmann IC, Parra GJ, Lanyon JM, Seddon JM (2012). Fine-scale genetic population structure in a mobile marine mammal: inshore bottlenose dolphins in Moreton Bay, Australia. Mol Ecol 21: 4472–4485.

Baker CS, Steel D, Calambokidis J, Falcone E, González-Peral U, Barlow J, et al. (2013). Strong maternal fidelity and natal philopatry shape genetic structure in North Pacific humpback whales. Mar Ecol Prog Ser 494: 291–306.

Bernatchez L, Wellenreuther M, Araneda C, Ashton DT, Barth JMI, Beacham TD, et al. (2017). Harnessing the Power of Genomics to Secure the Future of Seafood. Trends Ecol Evol 32: 665–680.

Best PB (1979). Social Organization in Sperm Whales, Physeter macrocephalus. In: Winn HE, Olla BL (eds) Behavior of Marine Animals, Springer US: Boston, MA, pp 227–289.

Bohonak AJ (1999). Dispersal, Gene Flow, and Population Structure. Q Rev Biol 74: 21–45.

Buffalo V (2021). Quantifying the relationship between genetic diversity and population size suggests natural selection cannot explain Lewontin’s Paradox (G Sella, D Weigel, G Sella, and M Pennell, Eds.). eLife 10: e67509.

Burgess SC, Treml EA, Marshall DJ (2012). How do dispersal costs and habitat selection influence realized population connectivity? Ecology 93: 1378–1387.

Byers JE, Pringle JM (2006). Going against the flow: retention, range limits and invasions in advective environments. Mar Ecol Prog Ser 313: 27–41.

Chen S, Zhou Y, Chen Y, Gu J (2018). fastp: an ultra-fast all-in-one FASTQ preprocessor. Bioinformatics 34: i884–i890.

Cros A, Toonen RJ, Donahue MJ, Karl SA (2017). Connecting Palau’s marine protected areas: a population genetic approach to conservation. Coral Reefs 36: 735–748.

Danecek P, Auton A, Abecasis G, Albers CA, Banks E, DePristo MA, et al. (2011). The variant call format and VCFtools. Bioinformatics 27: 2156–2158.

Day J, Clark JA, Williamson JE, Brown C, Gillings M (2019). Population genetic analyses reveal female reproductive philopatry in the oviparous Port Jackson shark. Mar Freshw Res 70: 986–994.

Deepwater Horizon Natural Resource Damage Assessment Trustees (2016). Deepwater Horizon oil spill: Final Programmatic Damage Assessment and Restoration Plan and Final Programmatic Environmental Impact Statement.

Do C, Waples RS, Peel D, Macbeth GM, Tillett BJ, Ovenden JR (2014a). NeEstimator v2: re- implementation of software for the estimation of contemporary effective population size (N) from genetic data. Mol Ecol Resour 14: 209–214.

Do C, Waples RS, Peel D, Macbeth GM, Tillett BJ, Ovenden JR (2014b). NeEstimator v2: relJimplementation of software for the estimation of contemporary effective population size (*N_e_*) from genetic data. Mol Ecol Resour 14: 209–214.

Dray S, Dufour A-B (2007). The ade4 package: implementing the duality diagram for ecologists. J Stat Softw 22: 1–20.

Engelhaupt D, Rus Hoelzel A, Nicholson C, Frantzis A, Mesnick S, Gero S, et al. (2009). Female philopatry in coastal basins and male dispersion across the North Atlantic in a highly mobile marine species, the sperm whale (Physeter macrocephalus). Mol Ecol 18: 4193–4205.

Estabrook B, Ponirakis D, Clark C, Rice A (2016). Widespread spatial and temporal extent of anthropogenic noise across the northeastern Gulf of Mexico shelf ecosystem. Endanger Species Res 30: 267–282.

Fan G, Zhang Y, Liu X, Wang J, Sun Z, Sun S, et al. (2019). The first chromosome-level genome for a marine mammal as a resource to study ecology and evolution. Mol Ecol Resour 19: 944–956.

Frankham R, Bradshaw CJA, Brook BW (2014). Genetics in conservation management: Revised recommendations for the 50/500 rules, Red List criteria and population viability analyses. Biol Conserv 170: 56–63.

Franklin I (1980). Evolutionary change in small populations. Conserv Biol: 135–149.

Gallego-García N, Caballero S, Shaffer HB (2021). Are Genomic Updates of Well-Studied Species Worth the Investment for Conservation? A Case Study of the Critically Endangered Magdalena River Turtle. J Hered 112: 575–589.

Garrison E, Marth G (2012). Haplotype-based variant detection from short-read sequencing.

Goudet J (2005). HEIRFSTAT, a package for R to compute and test hierarchical *F* lJstatistics. Mol Ecol Notes 5: 184–186.

Grosberg R, Cunningham CW (2001). Genetic structure in the sea. Mar Community Ecol: 61– 84.

Guichoux E, Lagache L, Wagner S, Chaumeil P, Léger P, Lepais O, et al. (2011). Current trends in microsatellite genotyping. Mol Ecol Resour 11: 591–611.

Hanghøj K, Moltke I, Andersen PA, Manica A, Korneliussen TS (2019). Fast and accurate relatedness estimation from high-throughput sequencing data in the presence of inbreeding. GigaScience 8: giz034.

Hayes SA, Josephson E, Maze-Foley K, Rosel PE, McCordic J, Brossard A, et al. (2024). U.S. Atlantic and Gulf of Mexico Marine Mammal Stock Assessments 2023. NOAA Tech Memo NMFS-NE-321.

Hayes SA, Josephson E, Maze-Foley K, Rosel PE, Turek J, Byrd B, et al. (2021). US Atlantic and Gulf of Mexico Marine Mammal Stock Assessments 2020.

Hemstrom W, Jones M (2023). snpR: User friendly population genomics for SNP data sets with categorical metadata. Mol Ecol Resour 23: 962–973.

Hendry AP, Castric V, Kinnison MT, Quinn TP, Hendry A, Stearns S (2004). The evolution of philopatry and dispersal. Evol Illum Salmon Their Relat: 52–91.

Hodel RGJ, Segovia-Salcedo MC, Landis JB, Crowl AA, Sun M, Liu X, et al. (2016). The Report of My Death was an Exaggeration: A Review for Researchers Using Microsatellites in the 21st Century. Appl Plant Sci 4.

Jacquard A (1972). Genetic information given by a relative. Biometrics: 1101–1114.

Jacquard A (1974). The Genetic Structure of Populations, 1970th edn. Springer Science & Business Media: Berlin, Germany.

Jeffries DL, Copp GH, Lawson Handley L, Olsén KH, Sayer CD, Hänfling B (2016). Comparing RADseq and microsatellites to infer complex phylogeographic patterns, an empirical perspective in the Crucian carp, Carassius carassius, L. Mol Ecol 25: 2997–3018.

Jochens AD, Benoit-Bird K, Engelhaupt, D, Gordon J, Hu C, Jaquet N, et al. Sperm whale seismic study in the Gulf of Mexico: Synthesis report. U.S. Dept. of the Interior, Minerals Management Service, Gulf of Mexico OCS Region: New Orleans, LA.

Jombart T (2008). adegenet: a R package for the multivariate analysis of genetic markers. Bioinformatics 24: 1403–1405.

Kamvar ZN, Tabima JF, Grünwald NJ (2014). Poppr: an R package for genetic analysis of populations with clonal, partially clonal, and/or sexual reproduction. PeerJ 2: e281.

Kelly RP, Palumbi SR (2010). Genetic Structure Among 50 Species of the Northeastern Pacific Rocky Intertidal Community. PLOS ONE 5: e8594.

Kocher TD, Thomas WK, Meyer A, Edwards SV, Pääbo S, Villablanca FX, et al. (1989). Dynamics of mitochondrial DNA evolution in animals: amplification and sequencing with conserved primers. Proc Natl Acad Sci 86: 6196–6200.

Komoroske LM, Jensen MP, Stewart KR, Shamblin BM, Dutton PH (2017). Advances in the Application of Genetics in Marine Turtle Biology and Conservation. Front Mar Sci 4.

Konrad CM, Gero S, Frasier T, Whitehead H (2018). Kinship influences sperm whale social organization within, but generally not among, social units. R Soc Open Sci 5: 180914.

Korunes KL, Samuk K (2021). pixylJ: Unbiased estimation of nucleotide diversity and divergence in the presence of missing data. Mol Ecol Resour 21: 1359–1368.

Lande R (1995). Mutation and Conservation. Conserv Biol 9: 782–791.

Largier JL (2003). Considerations in Estimating Larval Dispersal Distances from Oceanographic Data. Ecol Appl 13: 71–89.

Lavery TJ, Roudnew B, Gill P, Seymour J, Seuront L, Johnson G, et al. (2010). Iron defecation by sperm whales stimulates carbon export in the Southern Ocean. Proc R Soc B Biol Sci 277: 3527–3531.

Leigh JW, Bryant D, Nakagawa S (2015). POPART: full-feature software for haplotype network construction. Methods Ecol Evol 6.

Lemopoulos A, Prokkola JM, Uusi-Heikkilä S, Vasemägi A, Huusko A, Hyvärinen P, et al. (2019). Comparing RADseq and microsatellites for estimating genetic diversity and relatedness — Implications for brown trout conservation. Ecol Evol 9: 2106–2120.

Lizewski K, Alexander A, Steel D, Mate B, Engelhaupt D, Wise J, et al. (2025). Change in matrilineal structure over time in an isolated population of sperm whales. Mar Ecol Prog Ser 758: 161–178.

Mamoozadeh NR, Graves JE, McDowell JR (2020). Genome-wide SNPs resolve spatiotemporal patterns of connectivity within striped marlin (Kajikia audax), a broadly distributed and highly migratory pelagic species. Evol Appl 13: 677–698.

Manichaikul A, Mychaleckyj JC, Rich SS, Daly K, Sale M, Chen W-M (2010). Robust relationship inference in genome-wide association studies. Bioinformatics 26: 2867– 2873.

McKinney GJ, Waples RK, Seeb LW, Seeb JE (2017). Paralogs are revealed by proportion of heterozygotes and deviations in read ratios in genotypinglJbylJsequencing data from natural populations. Mol Ecol Resour 17: 656–669.

Mesnick SL, Taylor BL, Archer FI, Martien KK, Treviño SE, Hancock-Hanser BL, et al. (2011a). Sperm whale population structure in the eastern and central North Pacific inferred by the use of single-nucleotide polymorphisms, microsatellites and mitochondrial DNA. Mol Ecol Resour 11: 278–298.

Mesnick SL, Taylor BL, Archer FI, Martien KK, Treviño SE, Hancock-Hanser BL, et al. (2011b). Sperm whale population structure in the eastern and central North Pacific inferred by the use of single-nucleotide polymorphisms, microsatellites and mitochondrial DNA. Mol Ecol Resour 11: 278–298.

Mijangos JL, Gruber B, Berry O, Pacioni C, Georges A (2022). dartR v2: An accessible genetic analysis platform for conservation, ecology and agriculture. Methods Ecol Evol 13: 2150–2158.

Morgan SG, Fisher JL, Miller SH, McAfee ST, Largier JL (2009). Nearshore larval retention in a region of strong upwelling and recruitment limitation. Ecology 90: 3489–3502.

Morin PA, Foote AD, Baker CS, Hancock-Hanser BL, Kaschner K, Mate BR, et al. (2018). Demography or selection on linked cultural traits or genes? Investigating the driver of low mtDNA diversity in the sperm whale using complementary mitochondrial and nuclear genome analyses. Mol Ecol 27: 2604–2619.

Moura AE, Kenny JG, Chaudhuri R, Hughes MA, Welch AJ, Reisinger RR, et al. (2014). Population genomics of the killer whale indicates ecotype evolution in sympatry involving both selection and drift. Mol Ecol 23: 5179.

Nazareno AG, Bemmels JB, Dick CW, Lohmann LG (2017). Minimum sample sizes for population genomics: an empirical study from an Amazonian plant species. Mol Ecol Resour 17: 1136–1147.

O’Corry-Crowe G, Suydam R, Quakenbush L, Potgieter B, Harwood L, Litovka D, et al. (2018). Migratory culture, population structure and stock identity in North Pacific beluga whales (Delphinapterus leucas). PLOS ONE 13: e0194201.

OrtegalJOrtiz JG, Engelhaupt D, Winsor M, Mate BR, Rus Hoelzel A (2012). Kinship of longlJterm associates in the highly social sperm whale. Mol Ecol 21: 732–744.

Palumbi SR (1994). Genetic Divergence, Reproductive Isolation, and Marine Speciation. Annu Rev Ecol Syst 25: 547–572.

Palumbi SR (2003). Population Genetics, Demographic Connectivity, and the Design of Marine Reserves. Ecol Appl 13: 146–158.

Pante E, Simon-Bouhet B (2013). marmap: A Package for Importing, Plotting and Analyzing Bathymetric and Topographic Data in R. PLOS ONE 8: e73051.

Paradis E (2010). pegas: an R package for population genetics with an integrated–modular approach. Bioinformatics 26: 419–420.

Pilot M, Dahlheim ME, Hoelzel AR (2010). Social cohesion among kin, gene flow without dispersal and the evolution of population genetic structure in the killer whale (Orcinus orca). J Evol Biol 23: 20–31.

Poelstra JW, Montero BK, Lüdemann J, Yang Z, Rakotondranary SJ, Hohenlohe P, et al. (2022). RADseq data reveal a lack of admixture in a mouse lemur contact zone contrary to previous microsatellite results. Proc R Soc B Biol Sci 289: 20220596.

R Core Team (2024). R: A language and environment for statistical computing, R Foundation for Statistical. Computing.

Rambaut A (2002). Se-Al v.2.0a11. http://tree.bio.ed.ac.uk/software/seal/.

Richard KR, Dillon MC, Whitehead H, Wright JM (1996). Patterns of kinship in groups of free- living sperm whales (Physeter macrocephalus) revealed by multiple molecular genetic analyses. Proc Natl Acad Sci 93: 8792–8795.

Rigby RA, Stasinopoulos DM (2005). Generalized additive models for location, scale and shape. J R Stat Soc Ser C Appl Stat 54: 507–554.

Rochette NC, RiveralJColón AG, Catchen JM (2019). Stacks 2: Analytical methods for pairedlJend sequencing improve RADseqlJbased population genomics. Mol Ecol 28: 4737–4754.

Rosel PE (2003). PCR-based sex determination in Odontocete cetaceans. Conserv Genet 4: 647–649.

Russello MA, Waterhouse MD, Etter PD, Johnson EA (2015). From promise to practice: pairing non-invasive sampling with genomics in conservation. PeerJ 3: e1106.

Sanford E, Kelly MW (2011). Local Adaptation in Marine Invertebrates. Annu Rev Mar Sci 3: 509–535.

Santiago E, Caballero A, Köpke C, Novo I (2024). Estimation of the contemporary effective population size from SNP data while accounting for mating structure. Mol Ecol Resour 24: e13890.

Sinclair C, Sinclair J, Zolman E, Martinez A, Balmer B, Barry K (2015). Remote biopsy sampling field procedures for cetaceans used during the natural resource damage assessment of the MSC252 Deepwater Horizon oil spill. NOAA technical memorandum; National Marine Fisheries Service; Southeast Fisheries Science Center.

Solsona-Berga A, Frasier K, Posdaljian N, Baumann-Pickering S, Wiggins S, Soldevilla M, et al. (2024). Accounting for sperm whale population demographics in density estimation using passive acoustic monitoring. Mar Ecol Prog Ser 746: 121–140.

Templeton AR, Crandall KA, Sing CF (1992). A cladistic analysis of phenotypic associations with haplotypes inferred from restriction endonuclease mapping and DNA sequence data. III. Cladogram estimation. Genetics 132: 619–633.

Thia JA (2023). Guidelines for standardizing the application of discriminant analysis of principal components to genotype data. Mol Ecol Resour 23: 523–538.

Vasimuddin M, Misra S, Li H, Aluru S (2019). Efficient architecture-aware acceleration of BWA- MEM for multicore systems. In: 2019 IEEE international parallel and distributed processing symposium (IPDPS), IEEE, pp 314–324.

Violi B, de Jong MJ, Frantzis A, Alexiadou P, Tardy C, Ody D, et al. (2023). Genomics reveals the role of admixture in the evolution of structure among sperm whale populations within the Mediterranean Sea. Mol Ecol 32: 2715–2731.

Vollmer NL, Viricel A, Wilcox L, Katherine Moore M, Rosel PE (2011). The occurrence of mtDNA heteroplasmy in multiple cetacean species. Curr Genet 57: 115–131.

Waples RK, Albrechtsen A, Moltke I (2019). Allele frequencylJfree inference of close familial relationships from genotypes or lowlJdepth sequencing data. Mol Ecol 28: 35–48.

Waples RS, Anderson EC (2017). Purging putative siblings from population genetic data sets: a cautionary view. Mol Ecol 26: 1211–1224.

Weir BS, Cockerham CC (1984). Estimating F-statistics for the analysis of population structure. evolution: 1358–1370.

Whitehead H (1993). The behaviour of mature male sperm whales on the Galápagos Islands breeding grounds. Can J Zool 71: 689–699.

Whitehead H (2018). Sperm Whale. In: Encyclopedia of Marine Mammals, Elsevier, pp 919– 925.

Whitehead H, Shin M (2022). Current global population size, post-whaling trend and historical trajectory of sperm whales. Sci Rep 12: 19468.

