## Supplemental Information for "Mitochondrial structure despite nuclear panmixia: sex-specific dispersal dictates population structure in sperm whales"

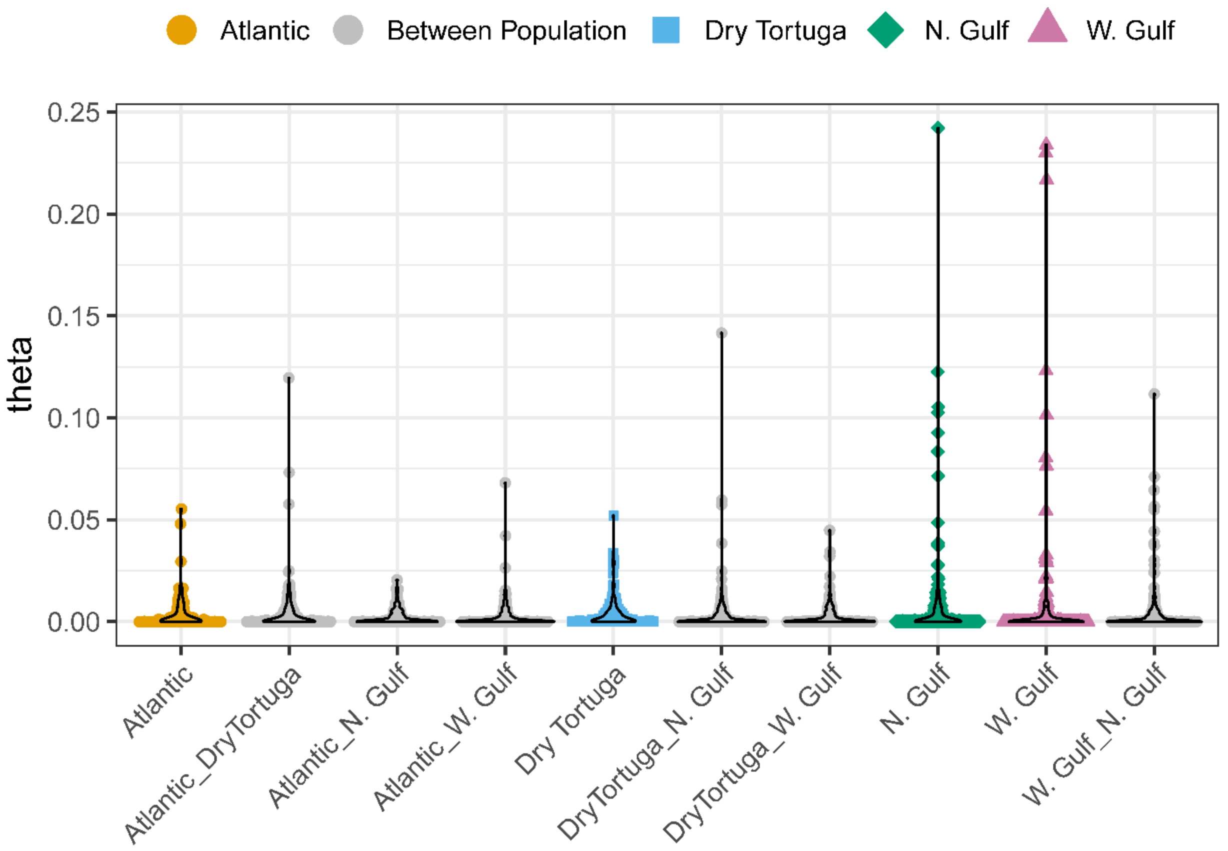


Figure S1: Pairwise relatedness categorized by population origin of each individual. Full siblings were found exclusively within geographic populations. See Figure 2 for geographic distance between individuals.

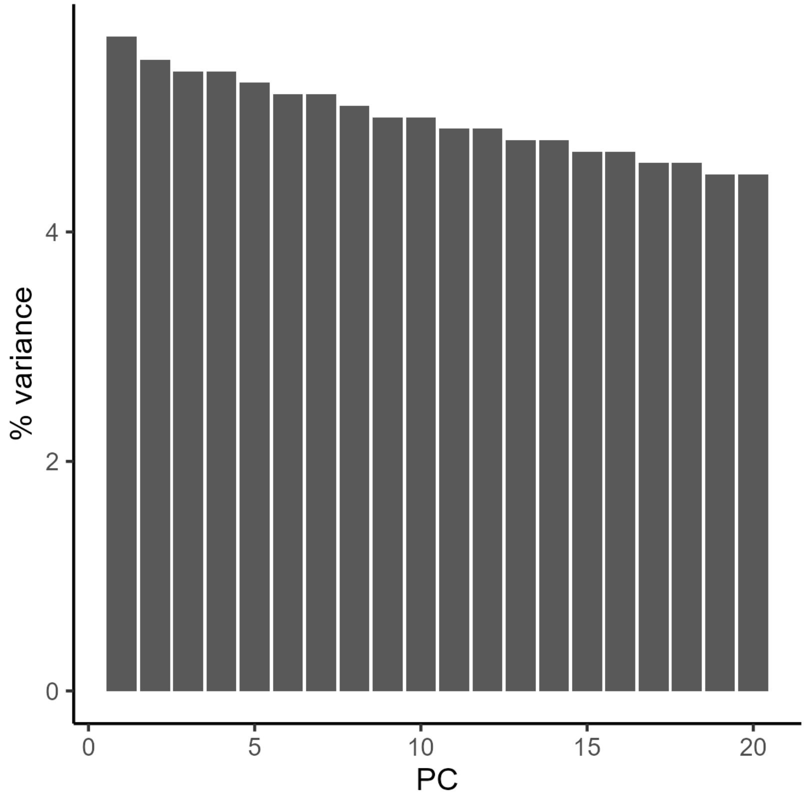


Figure S2: Percent variance explained by each principal component.

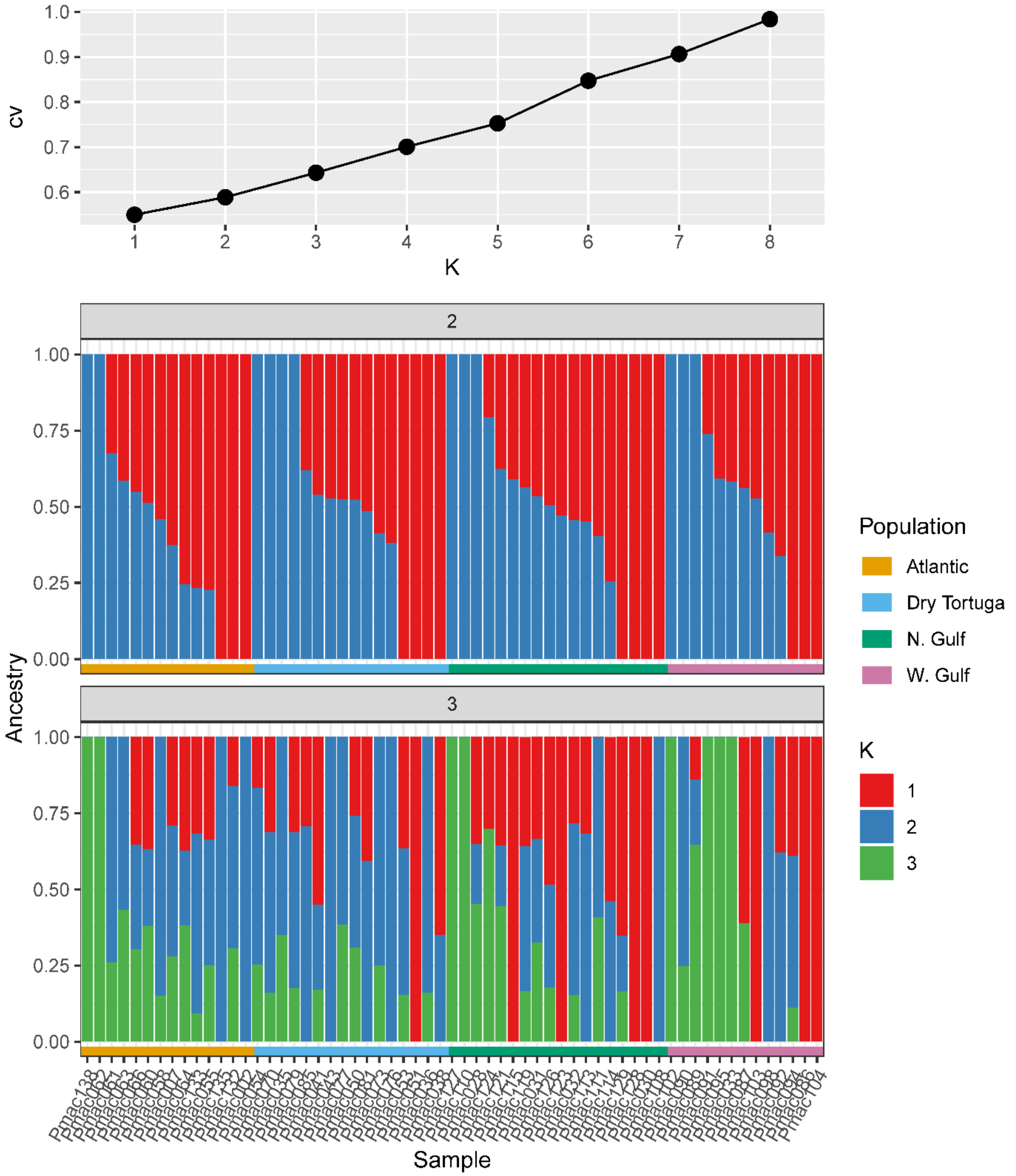


Figure S3: Admixture results for the genomic data. The top panel shows the cross validation error, where the most likely population can be inferred by the K with the lowest score. Here, the most likely number of populations is one. The next two panels show the estimated ancestry assuming two or threepopulations. Each bar is an individual where the color of the bar indicates the relative proportion of ancestry from each population. The horizontal bars below the plot indicate the population origin for each individual.

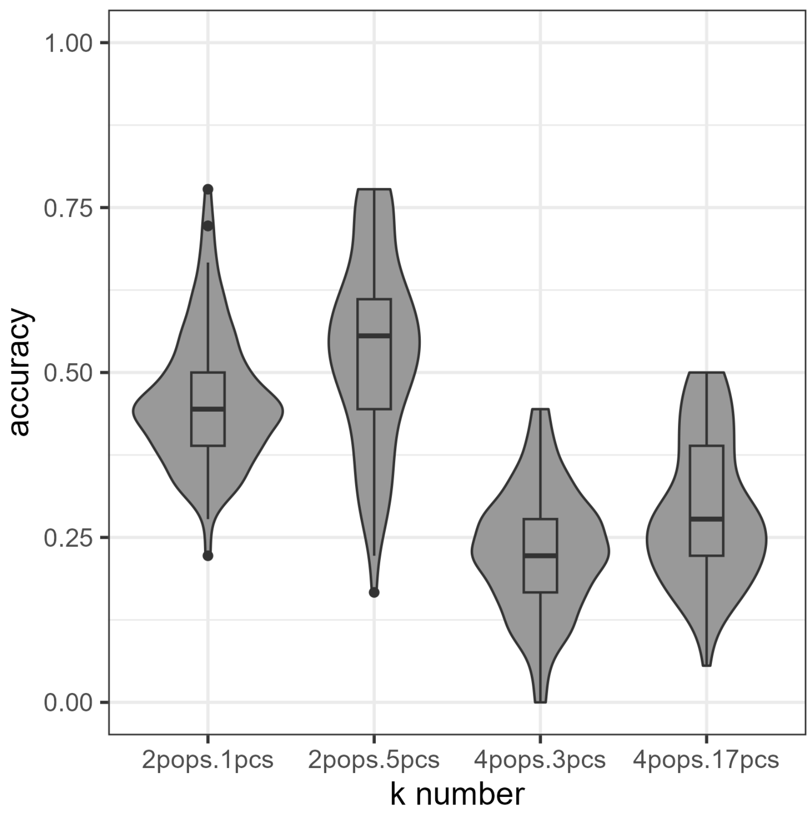


Figure S4: Assignment accuracy from DAPC following 100 replicates of training-testing partitioning assuming both 2 (Atlantic: Atlantic + Dry Tortuga; Gulf: N. Gulf + W. Gulf) and 4 populations and the number of principal components based on k-1 (2pops.1pcs; 4pops.3pcs) and the cross-validation approach (2pops.5pcs; 4pops.17pcs). If population assignments were completely random, we would expect 50% accuracy for the 2-population model and 25% accuracy for the 4-population model. The boxplot and violin plot show the distribution of mean accuracy for the 100 replicate runs.

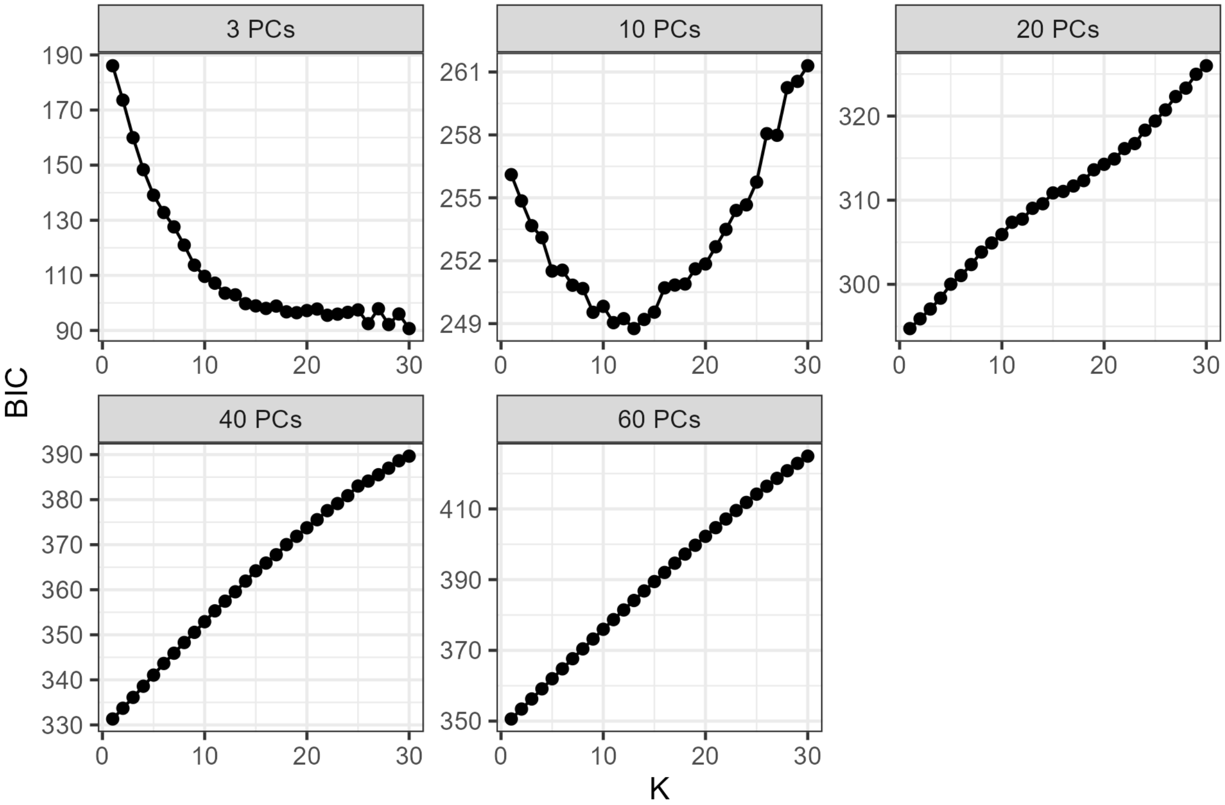


Figure S5: K-means clustering to determine the inferring number of populations. The x-axis shows the number of clusters (K) and the y-axis is the Bayesian information criterion (BIC) score. The K with the lowest BIC is typically considered the most likely number of populations. Note that these results are highly inconsistent with a changing number of PCs used, which suggests that population structure is highly limited or non-existent.

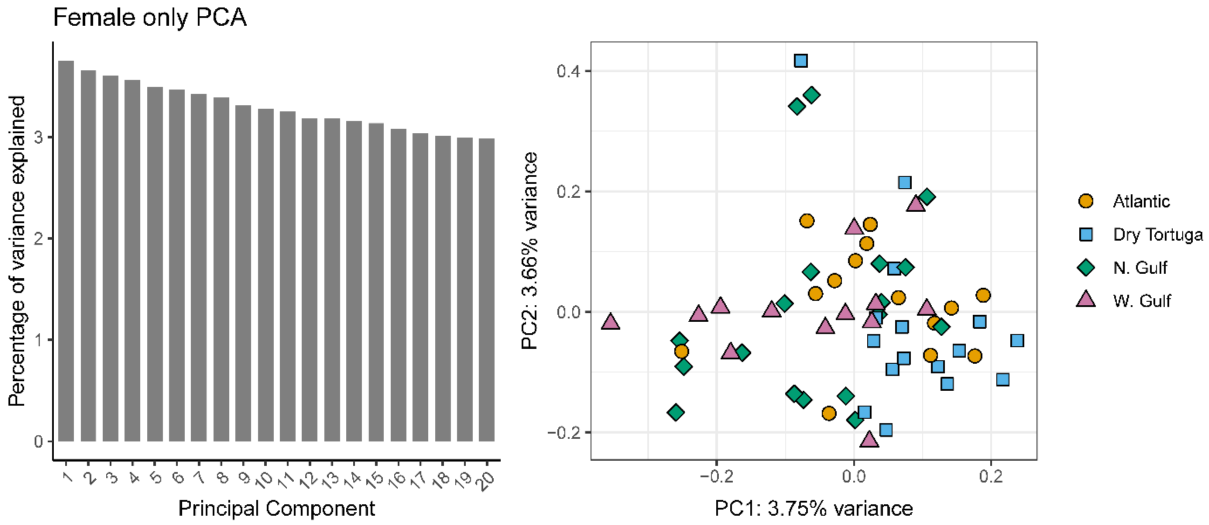


Figure S6: Subsetting the dataset to only female individuals, which show philopatry. Results are consistent with those found using the full dataset.

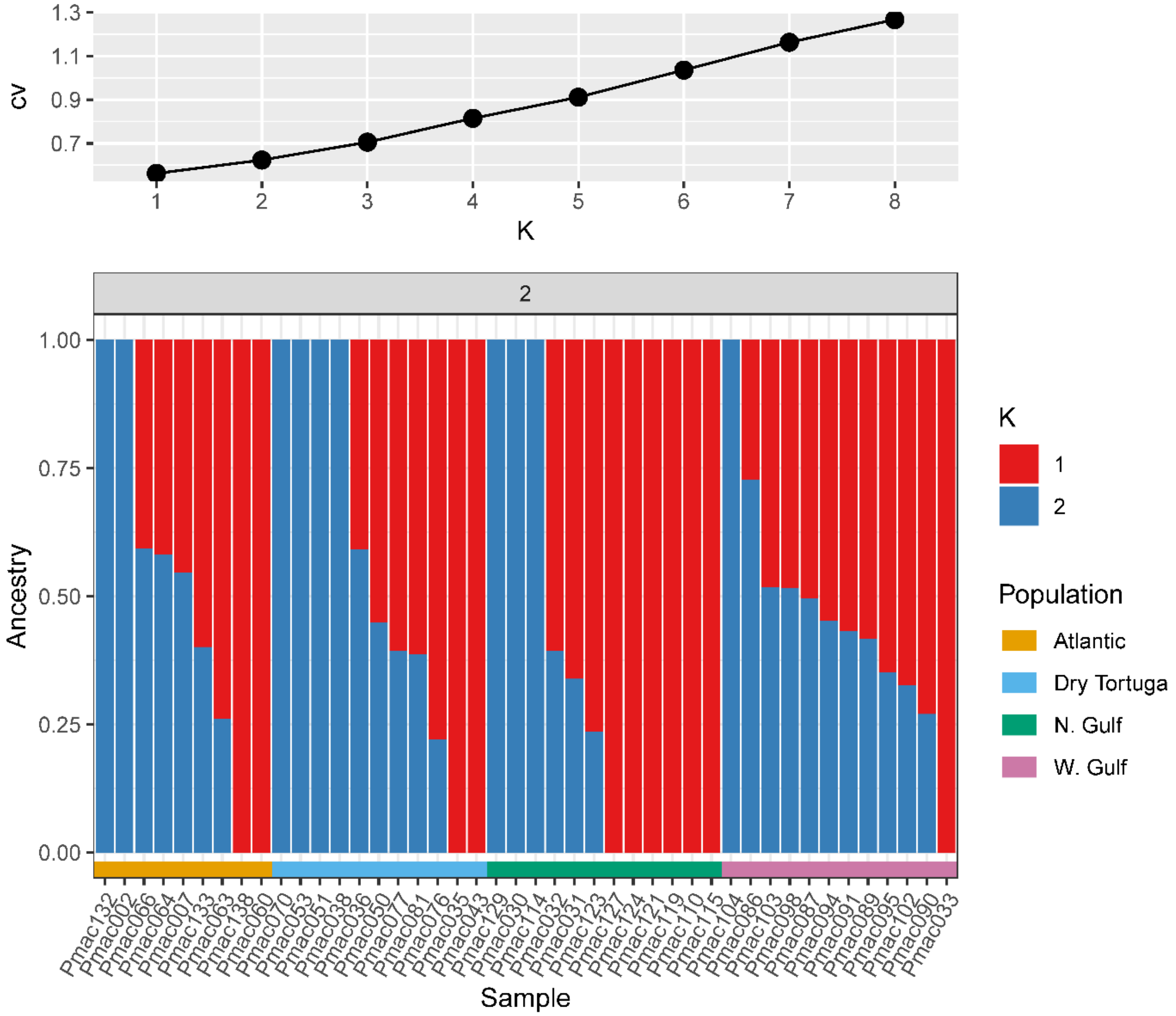


Fig. S7: Subsetting the genomic dataset to only female individuals, which show philopatry and testing for structure using admixture. Top panel shows the cross validation error, where the most likely population can be inferred by the K with the lowest score. Here, the most likely number of populations is one. The next panel shows the estimated ancestry assuming two populations. Each bar is an individual where the color of the bar indicates the relative proportion of ancestry from each population. The horizontal bars below the plot indicate the population origin for each individual. Note the consistency of these results with the full dataset.
